# *Lactobacillus* spp. decrease the susceptibility of *Salmonella* Typhimurium to the last resort antibiotic azithromycin

**DOI:** 10.1101/2023.10.01.560186

**Authors:** Lya Blais, Laurence Couture, Isabelle Laforest-Lapointe, Jean-Philippe Côté

## Abstract

Bacteria are involved in numerous interactions during infection and among host-associated microbial populations. *Salmonella enterica* serovar Typhimurium is a foodborne pathogen of great importance as well as a model organism to study interactions within a microbial community. In this study, we found that *S*. Typhimurium becomes tolerant to azithromycin when co-cultured with strains of *Lactobacillus*. Similarly, acidified media, from cell-free supernatant of *Lactobacillus* cultures for instance, also induced the tolerance of *S*. Typhimurium to azithromycin. The addition of membrane disruptors restored the normal sensitivity to azithromycin in acidified media, but not when *Lactobacillus* was present. These results suggested that the acidification of the media led to modification in envelope homeostasis, but that a different mechanism promoted the tolerance to azithromycin in the presence of *Lactobacillus* strains. To further understand how *Lactobacillus* strains modify the sensitivity of *S*. Typhimurium to azithromycin, a high-throughput assay was carried using the single gene deletion collection of the *S*. Typhimurium (1) in coculture with *L. rhamnosus* and (2) in sterile acidic conditions (pH 5.5 media only). As expected, both screens identified genes involved in envelope homeostasis and membrane permeability. Our results also suggest that changes in the metabolism of *S*. Typhimurium induce the tolerance observed in the presence of *L. rhamnosus*. Our results thus highlight two different mechanisms by which *Lactobacillus* strains induce tolerance of *S*. Typhimurium to antibiotics.

**Importance:** This study provides valuable insights into the intricate interactions between bacteria during infections and within host-associated microbial communities. Specifically, it sheds light on the significant role of *Lactobacillus* strains in inducing antibiotic tolerance in *Salmonella enterica* serovar Typhimurium, a critical foodborne pathogen and model organism for microbial community studies. The findings not only uncover the mechanisms underlying this antibiotic tolerance but also reveal two distinct pathways through which *Lactobacillus* strains might influence *Salmonella*’s response to antibiotics. Understanding these mechanisms has the potential to enhance our knowledge of bacterial infections and may have implications for the development of strategies to combat antibiotic resistance in pathogens like *Salmonella*. Furthermore, our results underscore the necessity to explore beyond the direct antimicrobial effects of antibiotics, emphasizing the broader microbial community context.

---

Bacterial pathogens are often found within polymicrobial communities and thus must interact with other microbes present in the environment. For instance, members of host-associated communities (*i*.*e*., microbiota) compete with pathogens entering a host and promote resistance to colonization by foreign microbes (1). Not surprisingly, pathogens have adapted to this polymicrobial environment and, for instance, overcome colonization resistance through the use of unique nutrients to persist in their ecological niche (2). There are many examples of the intricate relationships between pathogens and host-associated communities, but we are only beginning to understand the consequences of these interactions. A notorious example of interbacterial relationship is between *Pseudomonas aeruginosa* and *Staphylococcus aureus*, where *P. aeruginosa* profoundly alters the physiology of *S. aureus* and, accordingly, its susceptibility to antimicrobial agents (3–5). Therefore, investigating microbe-microbe interactions could provide new details about bacterial cells found in polymicrobial contexts. Herein, we probed the physiological state of the model intestinal pathogen *Salmonella enterica* serovar Typhimurium by determining its susceptibility against various antibiotics in the presence of *Lactobacillus* strains and found that *S*. Typhimurium becomes tolerant to azithromycin in presence of *Lactobacillus* isolates.

To explore the impact of *Lactobacillus* spp. on *S*. Typhimurium, we assessed the susceptibility of *S*. Typhimurium against various antibiotics during co-incubation experiments. Changes in antibiotic sensitivity are a powerful tool to assess the physiological state of the cell (6,7). We used selective media to determine the minimal inhibitory concentration (MIC) of antibiotics against *S*. Typhimurium co-incubated with one of two *Lactobacillus* strains, *L. rhamnosus* LMS2-1 or *L. reuteri* CF48-3A (Fig.S1A). A proof-of-concept experiments showed that, as expected, a beta-lactamase-producing *S. aureus* decreased the sensitivity of *S*. Typhimurium against ampicillin (Fig.S1B). We then assessed the susceptibility of *S*. Typhimurium against various antibiotics, including relevant antibiotics for treating salmonellosis such as ciprofloxacin, ceftriaxone and azithromycin (8,9), in the presence of either of the two *Lactobacillus* strains (Fig.1A). While we saw some isolate-specific effects, co-incubation with both *Lactobacillus* isolates resulted in a 16-fold increase in azithromycin MIC against *S*. Typhimurium (from 2 to 32 µg/mL; Fig.1A and Fig.S1C). Interestingly, *S*. Typhimurium seemed tolerant rather than resistant to azithromycin in the presence of the *Lactobacilli*, as cells recovered from the co-cultures where reverting back to their initial susceptibility to azithromycin (MIC of 2 µg/mL). In addition, heat-killed *Lactobacillus* cells did not increase the MIC to azithromycin against *S*. Typhimurium (Table S1).

**Figure 1.**
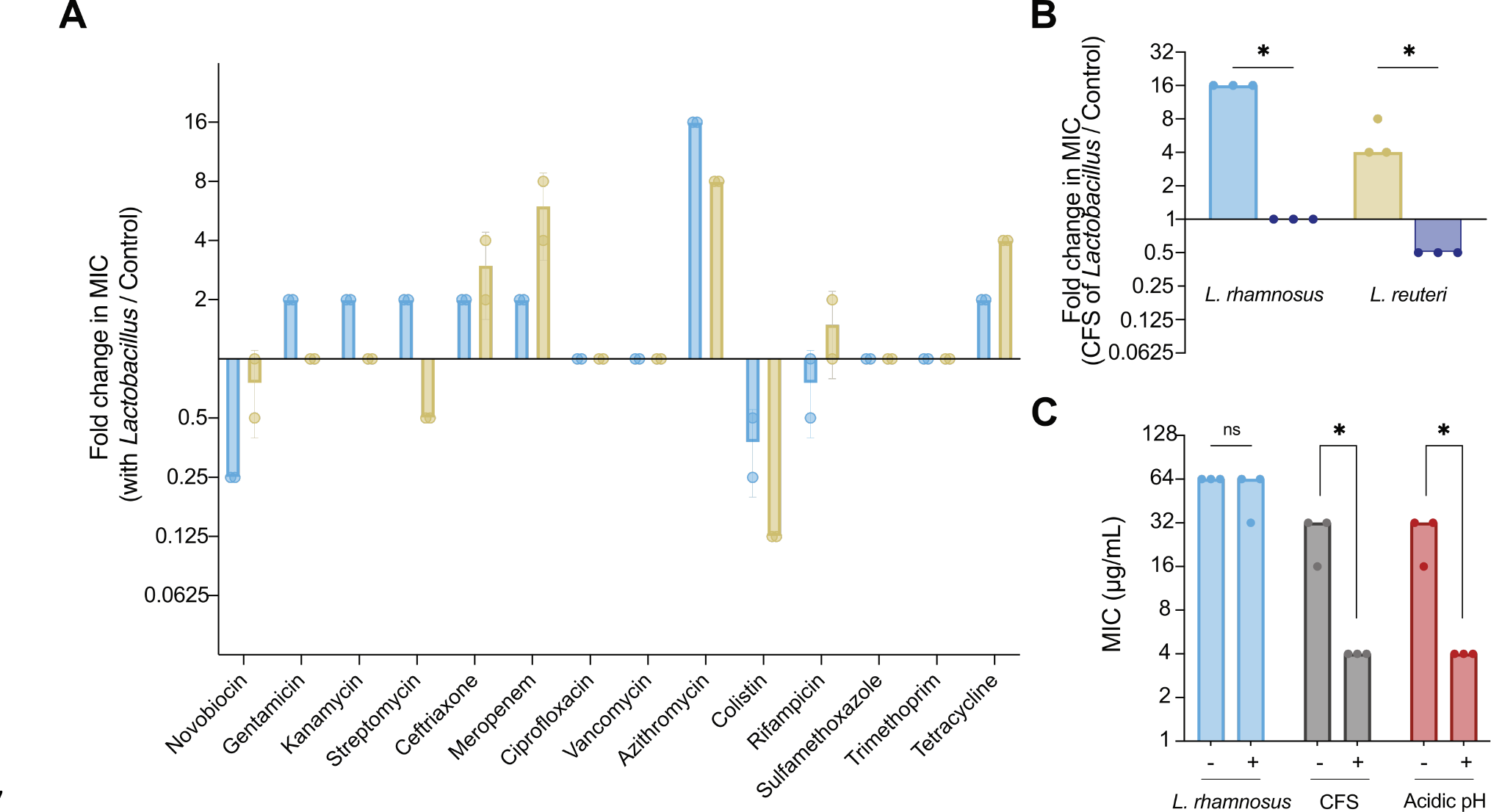
*Salmonella* Typhimurium becomes tolerant to azithromycin when cocultured with strains of *Lactobacillus*. (A) Fold change in the MIC of several antibiotics against *S*. Typhimuirum in the presence of *Lactobacillus* spp. Fold change was determined by producing a ratio of the MIC of *S*. Typhimurium in the presence of *L. rhamnosus* (blue) or *L. reuteri* (yellow) and the MIC of *S*. Typhimurium grown by itself against a selected panel of antibiotics. n=2 (B) Fold change in the MIC of azithromycin against *S*. Typhimurium in the presence of cell-free supernatant (CFS) (blue/yellow) or neutralized CFS (purple) of *Lactobacillus* spp. compared to the MIC of azithromycin against *S*. Typhimurium in monoculture. Initial pHs were 5.66 and 5.34 respectively for of *L. rhamnosus* and *L. reuteri* CFS. CFS was neutralized for both *Lactobacillus* to a pH of 7.00. Wilcoxon-Mann-Whitney U test, * indicates a p-value ≤ 0.05. n=3 (C) Impact of the membrane-active compound pentamidine on the MIC of azithromycin for *S*. Typhimurium in presence of *L. rhamnosus*, CFS or acidified media. Pentamidine was added at a concentration of 64 µg/mL. Wilcoxon-Mann-Whitney U test, * indicates a p-value ≤ 0.05. n=3.

Cell-free supernatants (CFS) from both *Lactobacillus* strains also induced azithromycin tolerance in *S*. Typhimurium (Fig.1B). As expected, CFS from the *Lactobacillus* strains were acidified compared to the media used for the experiment (pH of 5.66 for *L. rhamnosus* and 5.34 for *L. reuteri*). When neutralized CFS were used, the MICs went back to normal (Fig.1B). In addition, replacement of CFS by acidified media (BHI at pH 5.5) in combination with *S*. Typhimurium showed a similar increased in MIC (Table S1), suggesting that the decrease in pH caused the observed tolerance against azithromycin. In contrast, lactate, one of the main acidic compound produced by *Lactobacilli* (10), did not increase the MIC by itself (Table S1). It is important to note that azithromycin is stable in acidic conditions (9), and thus the tolerance to the antibiotic is not likely due to compound break down.

Azithromycin is dependent on envelope homeostasis to enter bacterial cells. Consequently, modification of cell permeability is a prevalent resistance mechanism (11,12). To investigate the hypothesis that the azithromycin tolerance was due to modified cell permeability and reduced azithromycin uptake, we conducted co-incubation assays in the presence of two membrane destabilizing agents: pentamidine and EDTA (13,14). The addition of pentamidine (Fig.1C) or EDTA (Fig.S2) still resulted in an increased MIC of azithromycin against *S*. Typhimurium when co-cultured with *L. rhamnosus*. However, when CFS of *L. rhamnosus* or acidified media were used, pentamidine and EDTA decreased the MIC of azithromycin compared to values observed for monocultures of *S*. Typhimurium. These results demonstrate the existence of two discrete and sperate mechanisms contributing to the observed phenotype. First, as expected, the acidification of the media most likely induced alterations in envelope permeability, decreasing the entry of azithromycin (15). Second, in the presence of *Lactobacillus*, a distinct mechanism emerges as a strong driver of the tolerance of *S*. Typhimurium to azithromycin.

To investigate the response of *S*. Typhimurium to the presence of *Lactobacillus* isolates or to acidified media, we reproduced the co-culture experiments to a larger scale, using a genome-wide single-gene deletion (SGD) collection of *S*. Typhimurium (16) and either high-density cultures of *L. rhamnosus* or acidified media, both in presence or absence of azithromycin. In our high-throughput experiments, the MIC of azithromycin against *S*. Typhimurium alone was 8 µg/mL. Therefore, we used a concentration of 128 µg/mL of azithromycin in our high-throughput assay to be representative of our initial observation (16-fold increase in MIC). At this concentration, *S*. Typhimurium only grows when *L. rhamnosus* is present or in acidified media, but not in monoculture. Mutants showing a growth defect specifically when treated with high concentrations of azithromycin indicate that their deleted gene is essential for the tolerance phenotype. We compared the growth of the SGD library with and without azithromycin in the presence of either (i) *L. rhamnosus* or (ii) acidified media. The two conditions showed different responses to the presence of azithromycin, with the presence of *L. rhamnosus* having a bigger impact on *S*. Typhimurium (Fig.2A), corroborating our hypothesis that different mechanisms are involved in the increased tolerance. We transformed screening values into Z-scores and selected gene deletions that were significatively different than the population (p-value ≤ 0.05) (Fig.2B and Table S3). While we did not necessarily observe the same hits, both screens were enriched for gene deletions known to disrupt the permeability of the outer membrane in *Escherichia coli* (*ldcA, mdoG, pal, pgm, prc, rfaP, rfaY, wecC, wecG, yceG* for instance) (17). In addition, in the presence of *L. rhamnosus*, the growth of both gene deletions from the two-component systems *envZ/ompR* and *phoP/phoQ* was impaired in the presence of azithromycin. The prevalence in genes affecting membrane homeostasis supports the effect of membrane disruptors in acidified media. In the presence of *L. rhamnosus*, additional gene deletions affecting core metabolic processes also impaired the tolerance of *S*. Typhimurium to azithromycin (*nuo* genes, *sdhAB, sucB, arcA* for instance). Our results suggests that, in the presence of *Lactobacillus, S*. Typhimurium undergoes changes in its metabolism leading to tolerance to azithromycin that prevails the modification in envelope homeostasis observed with acidified media.

**Figure 2.**
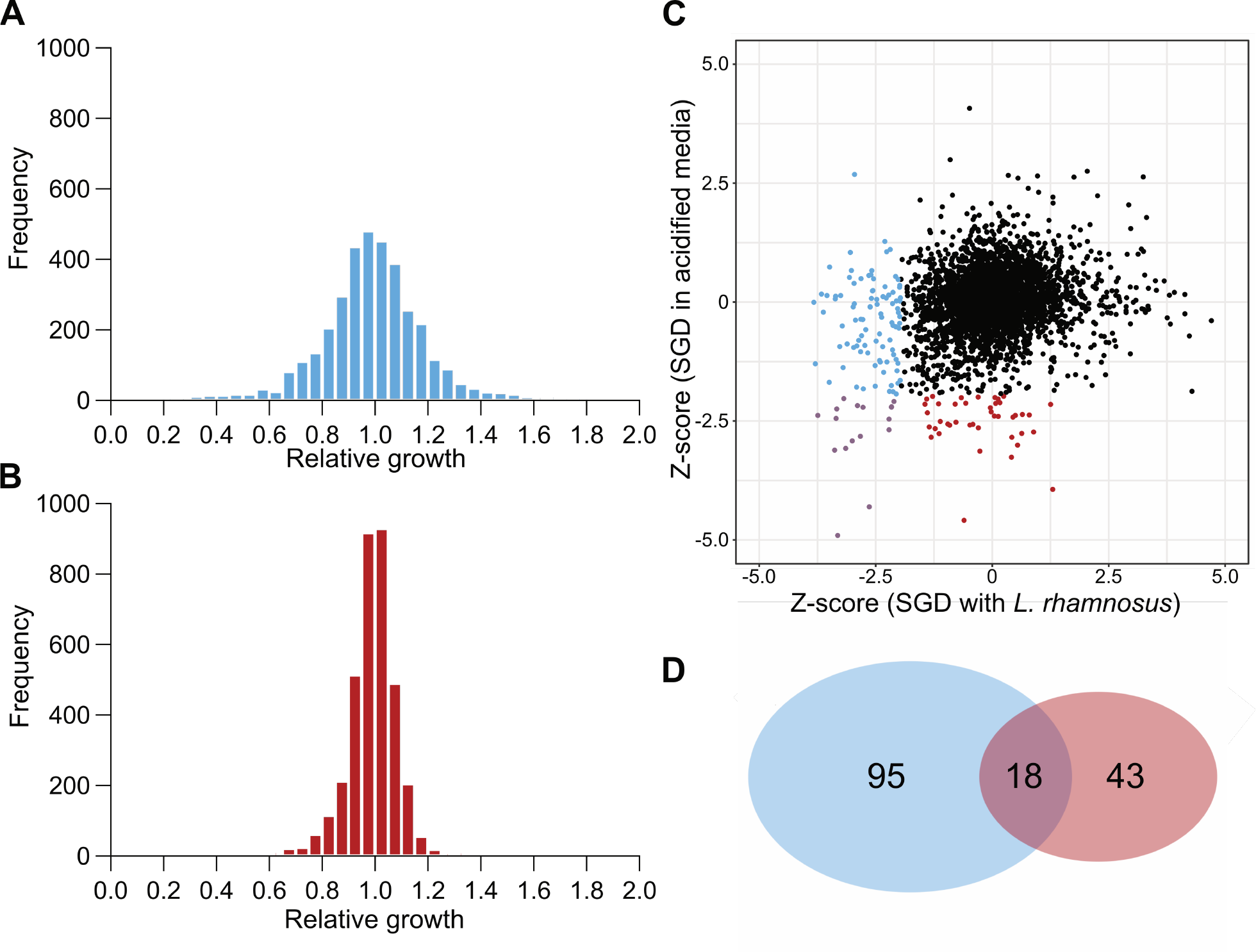
Growth of the SGD library at high azithromycin concentrations in the presence of *L. rhamnosus* or acidified pH. The SDG library was grown in the presence of (A) *L. rhamnosus* (blue) or (B) acidified media (red) and with 128 µg/mL of azithromycin (16X MIC in tested conditions). The growth of each mutant in the presence of azithromycin was compared to controls without azithromycin to yield a relative growth (a.u.) value, where a value of 1 means that azithromycin did not affect the growth of the mutant. (C) Relative growth values were converted to Z-scores. We selected hits corresponding to gene that were significatively different from the population (Z-score < -1.96, p-value ≤ 0.05) only the presence of *L. rhamnosus* (blue), acidified media (red) or in both conditions (purple). (D) Venn diagram of selected hits.

Further investigations of the interaction between *S*. Typhimurium and strains of *Lactobacillus* spp. will be necessary to fully understand the molecular basis behind how *Lactobacillus* isolates are able to induce a change in the susceptibility of the pathogen to azithromycin. If the enhanced tolerance of *S*. Typhimurium against azithromycin under acidic conditions is linked to modified membrane permeability, a distinct mechanism of action appears to be responsible for the same phenotype in presence of *L. rhamnosus*. In light of the escalating antibiotic resistance worldwide and the considerable obstacles to developing novel antibiotic molecules, our results underscore the urgency of delving deeper into the intricate dynamics between pathogens like *S*. Typhimurium and common human-associated bacteria like *Lactobacillus* spp., as they have the potential to significantly impact the effectiveness of antibiotic therapies.

## Acknowledgments

We thank all members of the JP Côté Lab at the University of Sherbrooke for their support and input on the project, and Daphnée Lamarche for critical review of the manuscript. This work was supported by the Natural Science and Engineering Research Council of Canada [NSERC; RGPIN-2019-06044] and by a starting grant for new investigators from the *Fonds de recherche du Québec – Santé* [FRQS; #295613]. J.P.C. holds a Chercheur boursier junior 1 fellowship from FRQS. I.L.-L. is supported by a Canada Research Chair T2. L.B. and L.C. were respectively supported by a master’s scholarship from the *Université de Sherbrooke* and a NSERC-USRA.

## Supplemental Material

### Materials and Methods

#### Bacterial strains and growth conditions

All experiments in this study were performed with *Salmonella enterica* serovar Typhimurium strain 14028s or the described single-gene deletion library (SGD library) of *S*. Typhimurium 14028s (16). *Lactobacillus reuteri* CF48-3A, *Lactobacillus rhamnosus* LMS2-1 and *Staphylococcus aureus* 130 were obtained from the Human Microbiome Project (18) and were cultured in Brain-Heart infusion (BHI) broth prepared according to Lau *et al* (19). All strains were grown at 37°C. The SGD library was maintained in BHI broth supplemented with kanamycin at a concentration of 50 mg/ml.

#### Products and solutions

A selective panel of antibiotic compounds was selected to cover various antibiotic classes. Azithromycin, meropenem and sulfamethoxazole were obtained from Sigma Aldrich and diluted in dimethyl sulfoxide (DMSO). Tetracycline, novobiocin, cefuroxime, ceftriaxone and colistin were also obtained from Sigma Aldrich but were diluted in water. Rifampicin, erythromycin, nalidixic acid and trimethoprim were obtained from Bio Basic and diluted in DMSO. Streptomycin, gentamicin, ampicillin, kanamycin, ciprofloxacin and vancomycin were also obtained from Bio Basic but were diluted in water. Lastly, apramycin was obtained from Alfa Aesar and was diluted in water. Ethylenediamine tetraacetic acid (EDTA) was obtained from Fisher Bioreagents and diluted in water. Pentamidine isethionate salt (Sigma Aldrich) was diluted in DMSO. All antibiotics and pentamidine were solubilized initially at 25mg/mL. Sodium L-lactate was also obtained from Sigma Aldrich and diluted in BHI.

#### Minimal inhibitory concentration assay

Determination of the minimal inhibitory concentration (MIC) was adapted from the Clinical and Laboratory Standards Institute (CLSI) guidelines (20). Briefly, each strain was grown overnight, then diluted 1:1000 in BHI. 2-fold dilution series of the antibiotics were added to the diluted culture in a 96-well plate starting at 256 µg/mL. The MIC, defined as the lowest concentration to inhibit visible growth, was determined at 18h incubation at 37°C. When indicated, EDTA or pentamidine was supplemented at a final concentration of 2 mM and 64 µg/mL, respectively. The MIC determination assay was also performed in acidic conditions as previously stated. Acidified BHI (pH of 5.5) was prepared with a solution of HCl 1N and used to dilute the different antibiotics to be tested in serial dilutions in 96-well plate.

#### Coculture minimal inhibitory concentration assay

The MIC of *S*. Typhimurium against various antibiotics was determined in the presence of *L. reuteri* CF48-3A or *L. rhamnosus* LMS2-1. The method for these combination assays was also adapted from the CLSI guidelines (20). Overnight cultures of *Lactobacillus* strains were grown for 18h at 37°C with agitation. *S*. Typhimurium was also grown overnight, then diluted 1:1000 in BHI. The co-culture experiments were performed by mixing the diluted culture of *Salmonella* with overnight cultures of the *Lactobacillus* strains in 96-well plates in a 1:1 ratio. Antibiotics were serially diluted as described for the MIC assays. The coculture plates were then incubated for 18h at 37°C. After the incubation, 5 µL of each well was plated on *Salmonella Shigella* (SS) agar, which is selective for Gram-negative bacteria. The agar plates were incubated overnight at 37°C, after which the MIC of *S*. Typhimurium was determined visually. The method is shown in Supplementary Figure S1A. When needed, EDTA or pentamidine were also added to the wells as described in the MIC assays.

For cocultures with the cell free supernatant (CFS) of *Lactobacillus* strains, *Lactobacillus* cultures were grown for 18h at 37°C with agitation. The overnight cultures were then centrifugated at 4,000 rpm for 10 min. The resulting supernatant was collected and filtered with a sterile 0.22 µm syringe filter. The sterile supernatants were then added to the 96-well plates containing the diluted *S*. Typhimurium and the antibiotics, as described above. For cocultures with heat-killed strains, cultures were grown for 18h at 37°C with agitation. After the incubation, strains were subjected to a heat treatment of 20 min at 70°C, followed by two rounds of 5 min centrifugation at 13,000 rpm and washing in 1X PBS. Washed pellet was resuspended in BHI before being added to the 96-well plates containing the diluted *S*. Typhimurium and the antibiotics, as described above. For the lactate assay, strains were grown as described above, and the lactate solution was added at a final concentration of 80 mM (21) in the 96-well plate containing the diluted *S*. Typhimurium and the antibiotics. It is important to note that the addition of the compound did not affect the initial pH of BHI, therefore the assay was conducted under neutral pH conditions.

#### High-throughput co-culture assay using the SGD collection

*S*. Typhimurium SGD collection was grown for 18h at 37°C in 384-well plates filled with 50 µL of BHI. *L. rhamnosus* LMS2-1 was grown separately in 25 ml of BHI at 37°C for 18h. The coculture plates were then prepared by adding 25 µL of overnight *L. rhamnosus* culture and 25 µL of BHI supplemented with 256 µg/mL of azithromycin in 384-well plates (final concentration of 128 µg/mL in 50 µl). The SGD collection was finally inoculated to the 384-well coculture plates with the Rotor HDA (Singer Instruments, UK). This setup transfers ~0.5 µl of the SGD culture into the new 384-well plates. The coculture plates were incubated for 18h at 37°C. The viability of *S*. Typhimurium was then determined by pinning from the 384-well coculture plates onto SS agar, which were incubated overnight at 37°C. Images of each plate were taken with the Phenobooth (Singer Instruments, UK) for analysis. This high-throughput method was also performed under acidic pH conditions. In this case, the 25 µL of the overnight culture of *L. rhamnosus* was replaced by 25 µL of BHI pH 5.5.

Images of the bacterial growth on SS agar plates were analyzed by an in-house analysis pipeline, as described previously (22). Briefly, imaged plates were divided into 384 region-of-interests (roi) to isolate each colony and the density within each roi was measured using Fiji (23). Density values were then normalized with the *lowess* in R (24). We determined to relative growth of every mutant of the collection for a given treatment (the presence of *L. rhamnosus* or acidified media) by calculation the ratio of the normalized density value with azithromycin and without azithromycin. A relative growth around 1 signifies that the gene deletion growth is similar with or without azithromycin, while a relative growth < 1 means that the growth of gene deletion is impaired in the presence of azithromycin. Relative growth values were then converted to Z-scores, Z score = (Conjugation score – screen mean)/screen sd. Based on the statistical distribution obtained from the Z-scores, we selected the gene deletions that were the most distant from the dataset average (p-values ≤0.05).

## Figures & Tables

**Figure S1.**
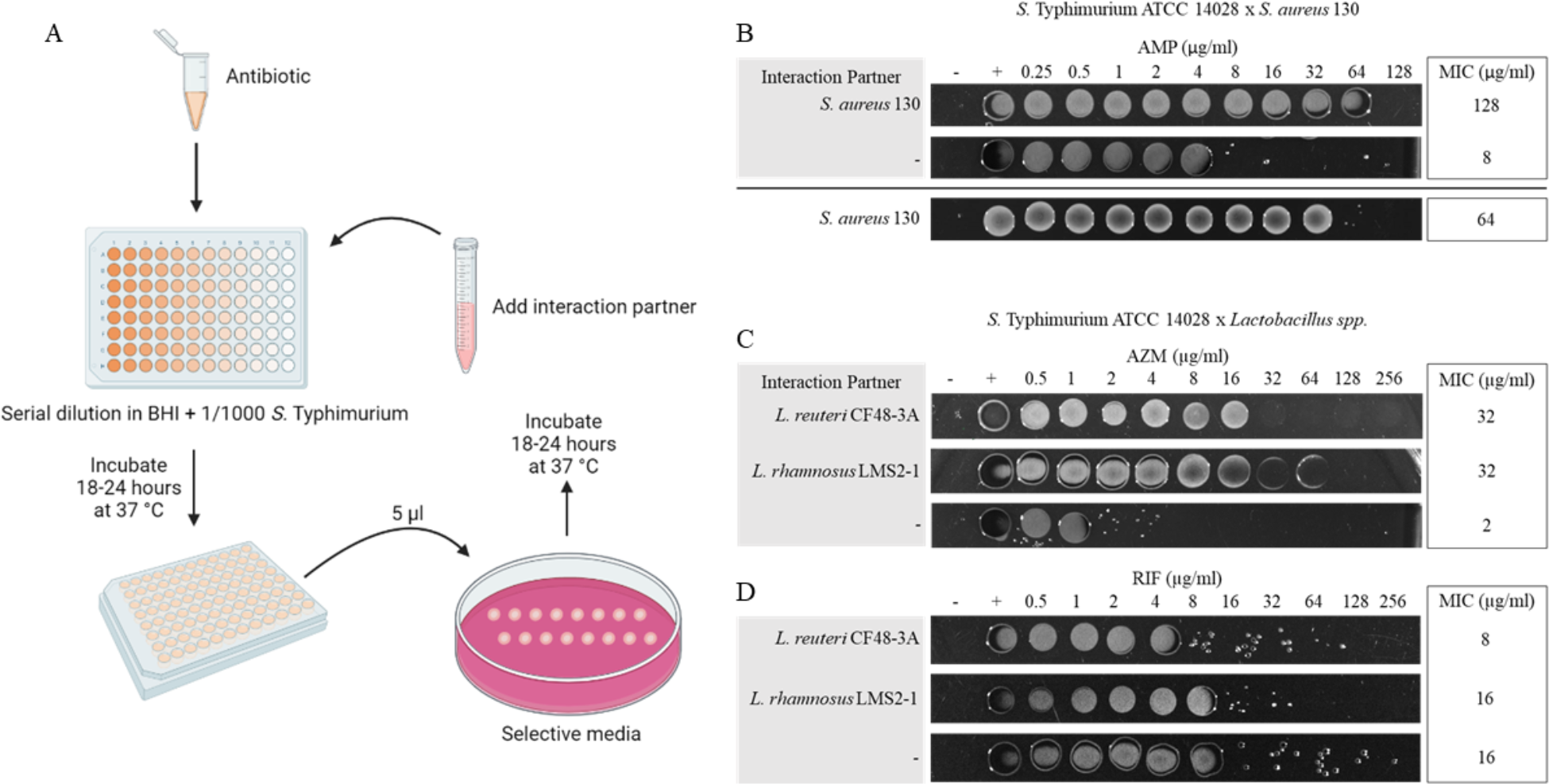
Coculture minimal inhibitory concentration assay. (A) Summary of the method used to determine the susceptibility of *S*. Typhimurium to different antibiotics following a co-culture. Co-cultures were performed in BHI media. After incubation, *S*. Typhimurium is selected by spotting 5 µL of the co-cultures on SS agar. (B) Proof-of-concept interaction between *S*. Typhimurium and *S. aureus* 130. The combination was done in the presence of increasing concentrations of ampicillin. MIC against *S*. Typhimurium is shown on SS agar. MIC against *S. aureus* is shown on LB agar. (C) Changes in the susceptibility of *S*. Typhimurium to azithromycin when cocultured with *Lactobacillus* spp. MIC against *S*. Typhimurium is shown on SS agar either alone with the antibiotic or in combination with *Lactobacillus*. (D) Strain combinations performed with rifampicin which is not impacted by the co-culture.

**Figure S2.**
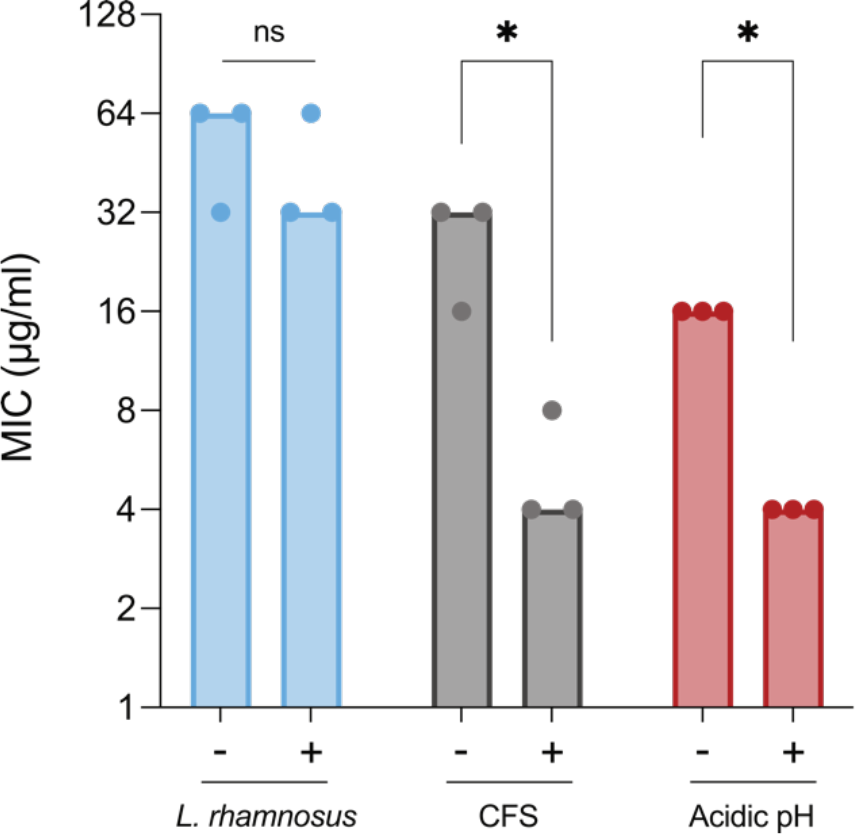
Addition of membrane-destabilizing agents in coculture MIC assays. Impact of EDTA on the MIC of azithromycin against *S*. Typhimurium in presence of *L. rhamnosus* culture, *L. rhamnosus* derived CFS or an acidified media. EDTA was added at a concentration of 2 mM. Wilcoxon-Mann-Whitney U test, * indicates a p-value ≤ 0.05.

**Table S1.** MIC of azithromycin against *S*. Typhimurium in various conditions.

| Strain | Combination | MIC of azithromycin (µg/mL) |
| --- | --- | --- |
| *S. Typhimurium* | - | 2-4 |
|  | *L. rhamnosus* | 32-64 |
|  | *L. rhamnosus* CFS | 32-64 |
|  | *L. rhamnosus* neutralized CFS | 4-8 |
|  | *L. reuteri* | 32-64 |
|  | *L. reuteri* CFS | 32-64 |
|  | *L. reuteri* neutralized CFS | 2-4 |
|  | BHI (pH 5.5) | 32-64 |
|  | 80mM lactate | 2 |
|  | Heat-killed *L. rhamnosus* | 2 |

**Table S2.** MIC of azithromycin against *S*. Typhimurium in different combination with the addition of membrane-destabilizing agents.

| Strain | Combination | MIC of azithromycin (µg/mL) |  |  |
| --- | --- | --- | --- | --- |
|  |  |  | + EDTA | + pentamidine |
| *S. Typhimurium* | - | 2 | 0.5 | 0.5 |
|  | *L. rhamnosus* | 32-64 | 32-64 | 32-64 |
|  | *L. reuteri* | 16-64 | 32-64 | 32-64 |
|  | BHI (pH 5.5) | 16-32 | 4 | 4 |
|  | *L. rhamnosus* CFS | 16-32 | 4-8 | 4 |

Note: EDTA was used at 2 mM and pentamidine at 64 µg/mL.

**Table S3. Relative growth and Z-score values for the complete SGD collection in the presence of *L. rhamnosus* or in acidified conditions as well as for selected hits.** See Excel file.

## References

1. Bäumler AJ, Sperandio V. Interactions between the microbiota and pathogenic bacteria in the gut. Nature. 2016 Jul 6;535(7610):85–93.

2. Sorbara MT, Pamer EG. Interbacterial mechanisms of colonization resistance and the strategies pathogens use to overcome them. Mucosal Immunol. 2019 Jan;12(1):1–9.

3. Ibberson CB, Stacy A, Fleming D, Dees JL, Rumbaugh K, Gilmore MS, et al. Coinfecting microorganisms dramatically alter pathogen gene essentiality during polymicrobial infection. Nat Microbiol. 2017 May;2:17079.

4. Beaudoin T, Yau YCW, Stapleton PJ, Gong Y, Wang PW, Guttman DS, et al. Staphylococcus aureus interaction with Pseudomonas aeruginosa biofilm enhances tobramycin resistance. npj Biofilms Microbiomes. 2017 Oct 19;3(1):1–9.

5. Orazi G, Jean-Pierre F, O’Toole GA. Pseudomonas aeruginosa PA14 Enhances the Efficacy of Norfloxacin against Staphylococcus aureus Newman Biofilms. J Bacteriol. 2020 Aug 25;202(18):e00159–20.

6. Hogan AM, Cardona ST. Gradients in gene essentiality reshape antibacterial research. FEMS Microbiol Rev. 2022 Feb 1;46(3):fuac005.

7. Farha MA, French S, Brown ED. Systems-Level Chemical Biology to Accelerate Antibiotic Drug Discovery. Acc Chem Res. 2021 Apr 20;54(8):1909–20.

8. Acheson D, Hohmann EL. Nontyphoidal Salmonellosis. Clinical Infectious Diseases. 2001 Jan 15;32(2):263–9.

9. Anandabaskar N. Protein Synthesis Inhibitors. In: Paul A, Anandabaskar N, Mathaiyan J, Raj GM, editors. Introduction to Basics of Pharmacology and Toxicology: Volume 2 : Essentials of Systemic Pharmacology : From Principles to Practice. Springer Nature; 2021. p. 835–68.

10. De Keersmaecker SCJ, Verhoeven TLA, Desair J, Marchal K, Vanderleyden J, Nagy I. Strong antimicrobial activity of Lactobacillus rhamnosus GG against Salmonella typhimurium is due to accumulation of lactic acid. FEMS Microbiol Lett. 2006 Jun;259(1):89–96.

11. Gomes C, Martínez-Puchol S, Palma N, Horna G, Ruiz-Roldán L, Pons MJ, et al. Macrolide resistance mechanisms in Enterobacteriaceae: Focus on azithromycin. Critical Reviews in Microbiology. 2017 Jan 2;43(1):1–30.

12. Farha MA, French S, Stokes JM, Brown ED. Bicarbonate Alters Bacterial Susceptibility to Antibiotics by Targeting the Proton Motive Force. ACS infectious diseases. 2018 Mar;4(3):382–90.

13. Stokes JM, MacNair CR, Ilyas B, French S, Côté JP, Bouwman C, et al. Pentamidine sensitizes Gram-negative pathogens to antibiotics and overcomes acquired colistin resistance. Nat Microbiol. 2017 Mar 6;2:17028.

14. MacNair CR, Brown ED. Outer Membrane Disruption Overcomes Intrinsic, Acquired, and Spontaneous Antibiotic Resistance. mBio. 2020 Sep 22;11(5):e01615–20.

15. Laub R, Schneider YJ, Trouet A. Antibiotic Susceptibility of Salmonella spp. at Different pH Values. Microbiology. 1989 Jun 1;135(6):1407–16.

16. Porwollik S, Santiviago CA, Cheng P, Long F, Desai P, Fredlund J, et al. Defined Single-Gene and Multi-Gene Deletion Mutant Collections in Salmonella enterica sv Typhimurium. PLoS ONE. 2014 Jul;9(7):e99820.

17. Stokes JM, French S, Ovchinnikova OG, Bouwman C, Whitfield C, Brown ED. Cold Stress Makes Escherichia coli Susceptible to Glycopeptide Antibiotics by Altering Outer Membrane Integrity. Cell Chemical Biology. 2016 Feb;23(2):267–77.

18. Turnbaugh PJ, Ley RE, Hamady M, Fraser-Liggett CM, Knight R, Gordon JI. The human microbiome project. Nature. 2007 Oct;449(7164):804–10.

19. Lau JT, Whelan FJ, Herath I, Lee CH, Collins SM, Bercik P, et al. Capturing the diversity of the human gut microbiota through culture-enriched molecular profiling. Genome Medicine. 2016 Jul 1;8(1):72.

20. Clinical & Laboratory Standards Institute. M100Ed33 | Performance Standards for Antimicrobial Susceptibility Testing, 33rd Edition.

21. Benito De Cárdenas IL, Ledesma OV, Pesce De Ruiz Holgado AA, Oliver G. Effect of Lactate on the Growth and Production of Diacetyl and Acetoin by Lactobacilli. Journal of Dairy Science. 1985 Aug;68(8):1897–901.

22. French S, Mangat C, Bharat A, Côté JP, Mori H, Brown ED. A robust platform for chemical genomics in bacterial systems. Molecular biology of the cell. 2016 Mar;27(6):1015–25.

23. Schindelin J, Arganda-Carreras I, Frise E, Kaynig V, Longair M, Pietzsch T, et al. Fiji: an open-source platform for biological-image analysis. Nat Methods. 2012 Jul;9(7):676–82.

24. R Core Team (2021). R: A language and environment for statistical computing. R Foundation for Statistical Computing, Vienna, Austria. https://www.R-project.org/.

